# A One-Year In Vivo Study of a Braided Pure Molybdenum Stent: Favorable Biocompatibility and the Critical Challenge of Vascular Biomechanics

**DOI:** 10.64898/2026.11.22.766907

**Authors:** Fujunhui Zhang, Baiyou Lin, Tianyu Wei, Zhaoxu Zheng, Jinbiao Yao, Weiliang Jin, Shuhan Wang, Weiming Liu, Bingqin Cai, Youyuan Xiang, Xiangdong Liu, Hui Huang, Yufeng Zheng, Yisen Zhang, Ming Li, Xinjian Yang

**Affiliations:** Department of Interventional Neuroradiology, Beijing Neurosurgical Institute, Capital Medical University, Beijing, China; Department of Neurosurgery, Beijing Tiantan Hospital, Capital Medical University, Beijing, China; Department of Non-active Medical Devices Testing, Shenzhen Institute for Drug Control (Shenzhen Testing Center of Medical Devices), Shenzhen, China; Conlife Medical Technology (Shenzhen) Co., Ltd., Shenzhen, China; School of Materials Science and Engineering, Peking University, Beijing, China; China-America Institute of Neuroscience and and Beijing Institute of Geriatrics, Xuanwu Hospital, Capital Medical University, Beijing, China; Beijing Institute of Brain Disorders, Capital Medical University, Beijing, China

## Abstract

**Background:** This study aims to evaluate a newly developed biodegradable stent made of pure Molybdenum (Mo) for treating supra-aortic arterial stenosis.

**Methods:** A 30.48 μm braided Mo stent was developed, and its mechanical performance and degradation process were tested in vitro. The stent was then implanted into the arteries of Beagle dogs for a one-year observation period. The study compared these new stents against standard non-degradable stents to monitor safety, structural changes, and the degradation rate within a living organism.

**Results:** The Mo stents demonstrated excellent biocompatibility, showing no signs of organ damage or neurotoxicity, and the material degraded at a rate suitable for aneurysm healing (4.88–5.12 μm/year). However, while the control stents held their shape, the Mo stents showed progressive structural deformation and recoil over time when implanted in the common carotid artery and subclavian artery of Beagle dogs.

**Conclusion:** While this study confirms that Mo is a biocompatible material for stents, it highlights a major challenge in supra-aortic arteries. Future designs must focus on improving the stent’s resistance to withstand the repetitive forces found in supra-aortic arteries before it can be considered for clinical use, and it still requires validation using atherosclerotic stenosis models in the future.

## 1. INTRODUCTION

Atherosclerosis is a major cause of supra-aortic arterial stenosis, which may involve the common carotid and subclavian arteries[1]. Carotid artery stenosis is one of the leading causes of ischemic stroke worldwide, accounting for approximately 10%–15% of all ischemic strokes and contributing substantially to stroke-related disability and mortality[2–4]. Subclavian artery stenosis may cause upper-limb claudication, vertebrobasilar insufficiency, angina, and subclavian steal syndrome[5,6]. Stenting has been shown to confer clinical benefits in patients with carotid artery stenosis but may induce in-stent restenosis and chronic inflammation through mechanical irritation and foreign-body reactions[7–9]. To overcome these limitations, drug-eluting stents incorporating antiproliferative agents have been developed to reduce the incidence of in-stent restenosis[10]. Nevertheless, as permanent intravascular implants, drug-eluting stents remain associated with late thrombosis, restenosis, and chronic inflammation one or two years after implantation. Biodegradable stents have emerged as a promising alternative to permanent metallic stents by providing transient mechanical support during vascular healing while avoiding the long-term effects[11].

Current biodegradable stent materials primarily include magnesium-based alloys, iron-based alloys, and polymers such as poly-L-lactic acid (PLLA)[12]. These material systems, while extensively investigated, reflect inherent shortcomings that limit their clinical performance. Magnesium-based stents exhibit favorable biocompatibility and biodegradability[13], but rapid degradation, relatively weak mechanical properties, and poor radiographic visibility limit their long-term performance. The frequent reliance on thicker struts to compensate for these shortcomings may, in turn, increase the risks of neointimal hyperplasia, thrombosis, and reduced deliverability[14,15]. In contrast, PLLA stents allow tunable degradation via molecular design[12], yet their relatively low radial strength, poor radiographic visibility, and bio-inert surface properties heighten the risks of early recoil, lumen loss, and delayed endothelialization, elevating late thrombosis risk[16–18]. Iron-based stents offer excellent mechanical strength, but their slow and pitting degradation may compromise long-term structural integrity, result in persistent residues, and limit MRI compatibility[19,20]. Moreover, degradation products have been implicated in local inflammatory and hypersensitivity responses[19,20].

Molybdenum (Mo) has been found in previous studies to overcome the limitations of previous degradable materials, exhibiting excellent biocompatibility and demonstrating potential as a degradable stent material[14,21–24]. Specifically, its high elastic modulus, yield strength, and tensile strength enable the maintenance of adequate radial support at reduced strut thickness, while its relatively high density confers improved radiopacity compared with existing degradable metals[14,23]. From a biological perspective, Mo is an essential trace element involved in enzymatic processes, and its systemic handling—primarily via renal excretion—suggests a low risk of long-term accumulation[21,25]. These characteristics collectively position Mo as a promising material for intracranial stent applications. Nevertheless, its long-term degradation behavior and vascular response, particularly in the form of biodegradable braided stents, have not yet been fully elucidated.

In the present study, a biodegradable braided Mo stent with a wall thickness of 30.48 μm was developed. Its performance was systematically evaluated through a combination of in vitro mechanical characterization and degradation analyses, followed by a one-year in vivo assessment to examine long-term biocompatibility and structural evolution. For the in vivo investigation, Mo stents and structurally matched nickel–titanium (NiTi) inert controls were implanted into the bilateral arteries of healthy Beagle dogs, providing a physiologically relevant model that approximates human vascular conditions.

## 2. METHODS

### 2.1 Stent fabrication

Biodegradable braided Mo stents were manufactured from commercially pure Mo wire (purity ≥ 99.9%; wire diameter: 30.48 μm [0.0012 inch]; Fort Wayne Metals, IN, USA) using a 24-carrier horizontal braiding system (KBW-90-24-1; Xuzhou Henghui Braiding Machine Co., Ltd., Jiangsu, China). During braiding, each wire was set to a length of 50 mm. A 1-over-1 braiding configuration was applied, yielding a braid density of 50.8 picks per inch. The as-braided stents subsequently underwent vacuum heat treatment at 650°C for 30 minutes (RCF120; Kunshan ACXVAC Vacuum Equipment Co., Ltd., Jiangsu, China) to relieve internal stresses and stabilize the structure.

### 2.2 Stent morphology and elemental characterization

Surface morphology and microstructure features were characterized using scanning electron microscopy (SEM; FEI Quanta250FEG, Thermo Fisher Scientific Inc., OR, USA). Elemental composition was analyzed by energy-dispersive X-ray spectroscopy (EDS; Xflash SCU Gen 7, Bruker Nano Inc., Berlin, Germany). Prior to SEM observation, samples were sputter-coated with a thin gold layer to enhance surface conductivity and ensure high-resolution imaging.

### 2.3 Mechanical testing

Mechanical performance was assessed with particular attention to flexibility and vessel apposition, given their relevance to navigation and deployment in tortuous vasculature. Stents were deployed into custom-fabricated silicone channel models (inner diameter: 4 mm) incorporating both straight and curved segments, with a minimum curvature radius of 25 mm. Device conformability and wall apposition were documented photographically.

Radial compression testing was performed using a universal testing system (Instron 5944, Instron Inc., MA, USA) equipped with a dedicated radial compression fixture. Both Mo stents and NiTi control stents (nominal diameter: 4 mm) compressed from 4 mm to 0.7 mm at a constant crosshead speed of 10 mm/min, while force–displacement data were continuously recorded. Comparative analysis between groups at corresponding displacement points was performed using paired t-tests. All measurements were conducted in triplicate.

### 2.4 Potentiodynamic polarization testing

Electrochemical behavior was investigated in accordance with ASTM F2129-2025 standards to assess corrosion resistance under physiologically relevant conditions. Potentiodynamic polarization measurements were carried out using a three-electrode system (PARSTAT2273, AMETEK Inc., TN, USA) in phosphate-buffered saline (PBS, pH 7.4 ± 0.05) maintained at 37 ± 0.5 °C. The Mo wire served as the working electrode, with a saturated calomel electrode as the reference and a platinum mesh as the counter electrode. To minimize oxygen interference, the electrolyte was purged with nitrogen at a flow rate of 150 cm^2^/min for at least 30 minutes prior to testing, with continuous purging throughout the experiment. Following a 1-hour stabilization at open-circuit potential, polarization scans were performed at 1 mV/s. The breakdown potential (*E_b_*), defined as the potential at which current density exceeded 100 μA/cm², was recorded as an indicator of pitting susceptibility. Tests were repeated three times to ensure reproducibility.

### 2.5 In vitro degradation assessment

Immersion tests were conducted in PBS (pH 7.4 ± 0.05; 37 ± 0.5 °C) for up to 60 days, using a volume-to-surface-area ratio of 20 mL/cm² to approximate physiological conditions. At predetermined time points (days 1, 2, 4, 7, 14, 28, 42, 56, and 60), Mo stents (n = 5) were retrieved and gently rinsed with deionized water to remove residual electrolyte. To ensure accurate quantification of material loss, corrosion products were removed via ultrasonic cleaning in a chromic acid solution (200 g/L CrO3 + 10 g/L AgNO3) at 60°C for 5 minutes. This process was repeated until a constant mass was achieved. The cleaned samples were subsequently vacuum-dried and weighed using an analytical balance with a resolution of 0.01 mg (BP211D, Sartorius AG, Göttingen, Germany). The corrosion rate (CR, μm/year) was calculated according to ASTM G31-72 using the formula:

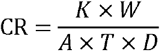

where K = 8.76 × 10□, W is mass loss (g), A is exposed surface area (cm²), T is immersion time (h), and D is the Mo density (10.28 g/cm³). The total surface area (A=1.15 cm²) was determined from the geometric configuration of the braided structure:

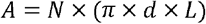

where N=24, d=30.48 μm, and L=50 mm.

Following 60 days of immersion, surface morphology was re-evaluated by SEM (Zeiss Sigma 360, Carl Zeiss Co., Inc., Baden-Württemberg, Germany) and elemental composition was analyzed using EDS (UltimMax40, Oxford Instruments plc., Oxfordshire, UK). Elemental maps and weight percentages (wt%) were recorded at baseline and 60 days to characterize compositional changes associated with degradation.

### 2.6 Animal study design and ethics

All animal procedures were approved by the Institutional Doctor (Guangzhou) Medical Technology Co., Ltd. (Approval No. IA024-1) and conducted in accordance with the Guide for the Care and Use of Laboratory Animals (8th edition, National Academies Press, 2011). Twenty adult male Beagle dogs (body weight: 10–15 kg; age: 18–24 months) were included in the study. Each animal received bilateral stent implantation: a Mo stent in one common carotid artery (CCA, n=10) and one subclavian artery (SCA, n=10), and a NiTi stent contralaterally. Animals were randomly assigned to sequential necropsies at 6 months, 9 months, and 12 months. All animals were maintained under controlled environmental conditions (12-hour light/dark cycle) with unrestricted access to standard chow and water throughout the study period.

### 2.7 Procedure details and peri-procedural management

To reduce thrombotic risk associated with endovascular intervention, dual antiplatelet therapy consisting of aspirin (50 mg) and clopidogrel (37.5 mg) was initiated three days prior to stent implantation and administered once daily. Food was withheld for 12 hours before the procedure. Under general anesthesia with endotracheal intubation, systemic anticoagulation was achieved using intravenous heparin (150 U/kg). The right femoral artery was surgically exposed and cannulated with a 5F vascular sheath. A 5F guiding catheter was advanced into the brachiocephalic trunk, and baseline digital subtraction angiography (DSA) was performed to measure the vessel diameter and confirm anatomical suitability. Stents were subsequently delivered to the target segments via a microcatheter and deployed under fluoroscopic guidance. Post-deployment DSA was performed immediately to verify accurate positioning and luminal patency. After device implantation, the sheath was removed, the femoral artery was ligated, and the incision was closed in anatomical layers. Postoperatively, animals were closely monitored twice daily for general condition and survival. Supportive care included a 5-day course of prophylactic antibiotics and analgesic administration for pain control. Dual antiplatelet therapy (aspirin 50 mg and clopidogrel 37.5 mg) was continued daily until the designated endpoint to minimize the risk of in-stent thrombosis.

### 2.8 In vivo imaging evaluation

Stent morphology was assessed using micro-computed tomography (micro-CT; SkyScan1172, Bruker Nano, Inc., Berlin, Germany) to provide high-resolution structural visualization. Longitudinal vascular assessment was performed using DSA (Azurion 3M15, Koninklijke Philips N.V., North Holland, The Netherlands) at baseline, 180 days, and 360 days, complemented by three-dimensional computed tomography angiography (3D-CTA; Incisive CT power, Koninklijke Philips N.V., North Holland, The Netherlands) at 90 days and Vascular Doppler ultrasonography (EPIQ 7C, Koninklijke Philips N.V., North Holland, The Netherlands) at 15 days. These imaging modalities were used to evaluate procedural integrity and detect adverse events, including thrombosis, in-stent stenosis (defined as ≥ 25% luminal diameter reduction), stent deformation, migration, and structural discontinuity. All images were independently reviewed by two blinded interventional neuroradiologists with more than 5 years of clinical experience. In cases of disagreement, the final adjudication was made by a senior physician with over 20 years of experience.

### 2.9 Laboratory tests

To characterize systemic exposure and potential toxicity, venous blood, spot urine, and fecal samples were collected under sedation at baseline and at 14, 30, 90, 180, and 360 days post-implantation. Mo concentrations in serum, urine, and fecal samples were quantified by inductively coupled plasma mass spectrometry ICP-MS (NexION 350X, PerkinElmer, Inc., MA, USA). Hematological and biochemical parameters were measured at 3, 14, 30, 90, 180, and 360 days to evaluate systemic responses. Results were interpreted against established canine reference ranges. Longitudinal comparisons with baseline values for each animal were performed using paired t-tests to detect statistically significant deviations.

### 2.10 Histological and immunohistochemical analysis

At 6 and 12 months, animals were euthanized by intravenous overdose of pentobarbital (100 mg/kg). Major organs, including the heart, liver, spleen, lungs, kidneys, and brain, were harvested, fixed in 10% neutral-buffered formalin for 48 hours, and processed for paraffin embedding. Hematoxylin and Eosin (H&E; BASO Diagnostics, Inc., Guangdong, China) staining was then performed. Brain regions (frontal, parietal, and occipital lobes) were further analyzed using immunohistochemistry with anti-GFAP (1:500; Novus Biologicals, Inc., CO, USA) for astrocytes, anti-IBA-1 (1:500; Guangzhou LBP Medicine Science & Technology Co., Ltd., Guangdong, China) for microglia, Luxol Fast Blue (LFB, Wuhan Servicebio Technology Co., Ltd., Hubei, China) for myelin, and Bielschowsky silver stain (Beijing Solarbio Science & Technology Co., Ltd., Beijing, China) for axonal integrity. Immunofluorescence was performed with DAPI (Wuhan Servicebio Technology Co., Ltd., Hubei, China) nuclear counterstaining and Fluoro-Jade C (FJC; Biosensis Pty Ltd., SA, Australia) for detection of degenerating neurons.

The stented vessel segments, including 5 mm margins proximal and distal to the stent, were harvested and fixed in 4% paraformaldehyde and subsequently dehydrated through a graded ethanol series. Following resin embedding, hard tissue sectioning was performed to obtain composite sections of the stent and vessel. The sections were subsequently stained with H&E and Masson’s trichrome for assessment of tissue morphology, collagen deposition, and smooth muscle cell proliferation. Whole-slide imaging was performed using a digital pathology slide scanner (Pannoramic SCAN, 3DHISTECH Ltd., Budapest, Hungary).

### 2.11 In vivo degradation characterization

In vivo degradation was quantified by measuring changes in the cross-sectional area of Mo wires using hard tissue sections obtained at 6, 9, and 12 months. Specimen preparation followed the protocol described in Section 2.10. Cross-sectional morphology of the stent–vessel complex was examined using SEM (FEI Quanta250FEG, Thermo Fisher Scientific Inc., OR, USA). For each node, four sections were analyzed, and five random sites per section were selected for quantitative measurement. The degradation rate and penetration depth were calculated based on cross-sectional area reduction:

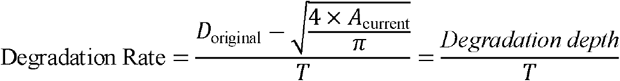

Where *D*_original_ represents the theoretical initial area of the Mo wire (30.48 μm), T represents implantation time (year) and *A*_current_ (μm^2^) represents the measured cross-sectional area derived from the SEM images using Image J (version 1.54r, National Institutes of Health, MD, USA). Longitudinal and cross-sectional morphology were examined by SEM/EDS as described in section 2.2

### 2.12 Statistical analysis

Continuous variables are presented as box plots, including median, interquartile range (IQR), and minimum–maximum values. Categorical variables are expressed as counts and percentages. Group comparisons for categorical data were compared using the chi-square test or Fisher’s exact test, as appropriate. Continuous variables, including laboratory measurements and mechanical testing outcomes, were analyzed using paired t-tests. Statistical analyses and graphical representations were generated using Graphpad Prism (version 10; GraphPad Software, LLC, CA, USA). A two-tailed *p* value < 0.05 was considered statistically significant.

## 3. RESULTS

### 3.1 Morphology and Mechanical Properties of the Stent

The braided Mo stent (4 × 20 mm) was fabricated using wires with a diameter of approximately 30.48 μm, as shown in Figure 1A. Figure 1B displays the surface topography and the corresponding EDS spectrum of the stent wire. Surface characterization (Figure 1B) revealed a smooth and homogeneous topography without evident defects, while EDS analysis confirmed the elemental composition to be predominantly pure Mo. Electrochemical evaluation further supported the material stability. The potentiodynamic polarization curve (Figure 1E) displayed a well-defined passive region, indicative of stable passive film formation on the stent surface. The measured breakdown potential (*E_b_*) reached 1840 mV, reflecting strong resistance to pitting corrosion and suggesting the capacity to maintain structural integrity under physiological conditions, an essential requirement for sustained mechanical support during aneurysm healing.

**Figure 1.**
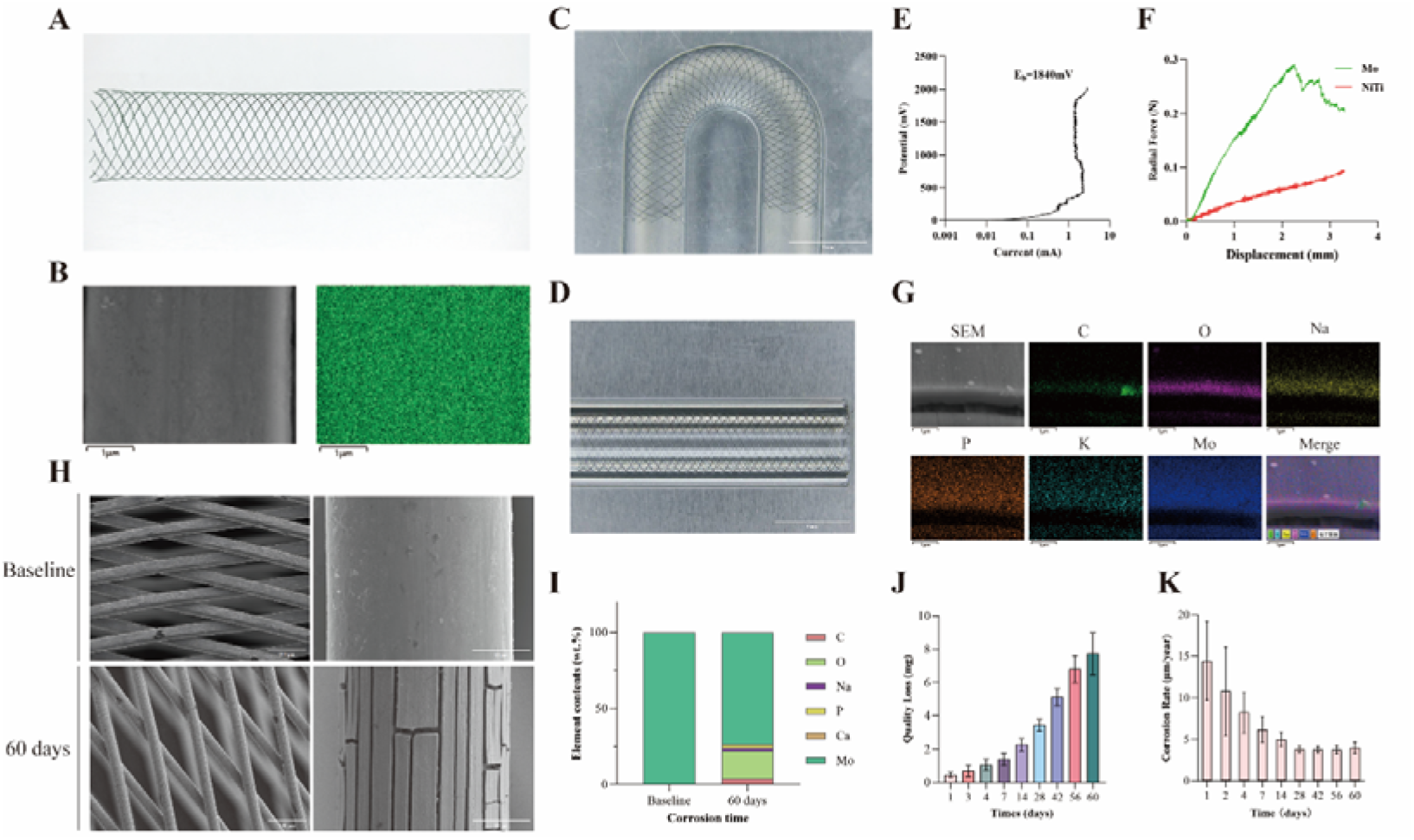
Characterization, mechanical behaviors, and in vitro degradation behaviors of the braided Mo stent. (A) Macroscopic view of the braided Mo stent. (B) Surface morphology and corresponding energy-dispersive spectroscopy (EDS) elemental mapping of the Mo wire. (C, D) Mechanical performance was evaluated by flexibility and apposition tests. (E) Potentiodynamic polarization curve of the Mo wire, showing a breakdown potential of 1840 mV. (F) Radial compression performance of the Mo stent compared with a NiTi stent of identical outer diameter (4 mm). (G-K) In vitro immersion tests. (G) Scanning electron microscope (SEM) image and EDS elemental maps after 60 days of immersion. (H) Surface morphology of the Mo stent after 60 days of immersion. (I) Elemental composition (wt%) of the Mo stent at baseline and after 60 days. (J) Mass loss of the Mo stent during immersion from 1 to 60 days. (K) Corrosion rate of the Mo stent over an immersion period of 1–60 days.

Mechanical performance testing demonstrated that the braided architecture provided both flexibility and conformability. In silicone vessel models (Figure 1C–D), the stent adapted closely to both straight and curved geometries without observable distortion, indicating favorable wall apposition even under simulated tortuous conditions. Radial compression testing (Figure 1F) revealed distinct mechanical behavior compared with NiTi controls. The Mo stent generated significantly higher radial force across the compression range (p < 0.001), with a gradual increase reaching a peak of 0.29 N (95% CI: 0.25– 0.33 N) at a compression diameter of 2.25 mm, followed by a decline. In contrast, the NiTi stent exhibited a more gradual and continuous increase in force, consistent with its superelastic characteristics. Collectively, these findings indicate that the Mo stent provides robust radial support while maintaining adequate flexibility.

### 3.2 In Vitro Degradation Behaviors

After 60 days of immersion, the stent surface was covered by a non-passivated degradation layer with widespread cracks (Figure 1H), reflecting progressive surface corrosion. Elemental analysis (Figure 1G, 1I) revealed that the layer primarily consists of C, O, Na, P, K, and Mo, with oxygen representing the dominant element aside from Mo, followed by C and P. This composition suggests that the degradation products are mainly Mo oxides with contributions from adsorbed ions in the simulated physiological environment.

Quantitative analysis of degradation kinetics demonstrated a time-dependent increase in mass loss over the 60 days (Figure 1J), consistent with ongoing material dissolution. The calculated corrosion rate (Figure 1K) exhibited a biphasic trend: a relatively rapid decrease during the early stage (days 1–14), likely reflecting initial surface activation and passive film formation, followed by stabilization at approximately 3.75–3.98 μm/year. This plateau indicates the establishment of a steady-state degradation process, suggesting a balance between material dissolution and surface film formation under in vitro conditions.

### 3.3 In vivo imaging evaluation of Mo and NiTi stents

In Figure 2, immediately after implantation, baseline DSA confirmed that both Mo and NiTi stents confirmed accurate deployment of both Mo and NiTi stents in the CCAs and SCAs, with satisfactory vessel wall apposition and preserved luminal patency. Longitudinal imaging revealed time-dependent structural changes in the Mo stents. At 90 days, CTA revealed localized distortion of the Mo stents in the SCAs, whereas the CCA stents remained intact. At 180 days, DSA showed progressive deformation of the CCA stents and re-coiling of the SCA stents, which became more pronounced by 360 days. Despite these structural alterations, no significant thrombosis or in-stent stenosis was detected at any stage, and the Mo stent maintained its overall integrity without significant degradation. Conversely, the NiTi control stents retained their original configuration and luminal patency throughout the 360-day observation period. Detailed structural changes and thrombosis events are summarized in Table 1. In the Mo group, the rates of stent thrombosis were 10.00% (2/20) at 15 days, and raised to 20.00% (4/20) at 90 days. No stent thrombosis was found in the NiTi group at any time point. The Mo group exhibited significantly higher rates of stent distortion than the NiTi group at 90, 180, and 360 days (50.00% vs. 5.00%, *P* = 0.001; 95.00% vs. 55.00%, *P* = 0.003; and 100.00% vs. 58.33%, *P* = 0.037, respectively). Additionally, stent recoil at 180 days was significantly more frequent in the Mo group than in the NiTi group (45.00% vs. 5.00%, P = 0.003), as was stent discontinuity at 360 days (46.15% vs. 0.00%, *P* = 0.015). No statistically significant between-group differences were observed for any other outcomes, including stent thrombosis, stent migration, and in-stent stenosis at any time point (all *P* > 0.05).

**Figure 2.**
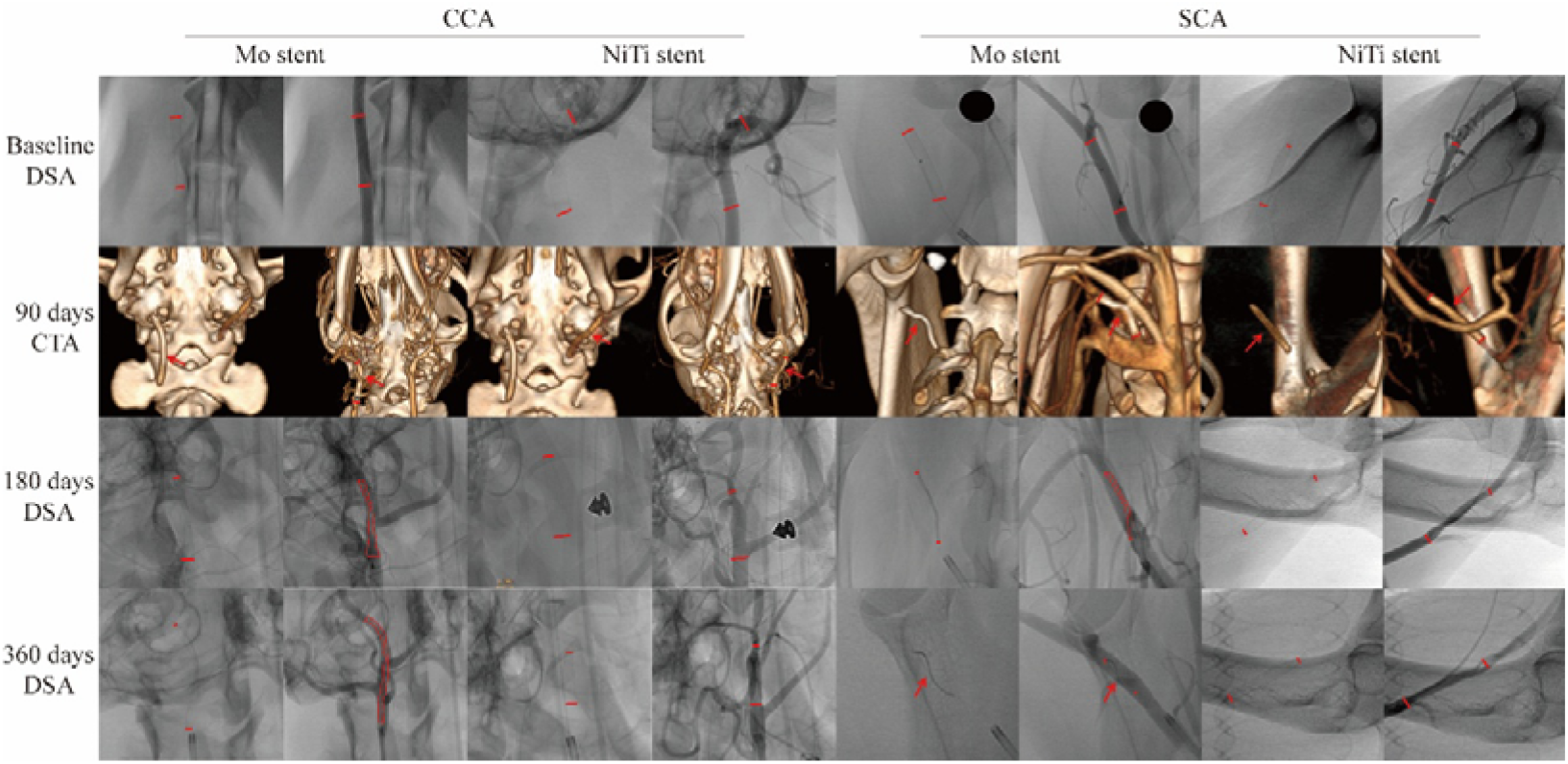
In vivo imaging of Mo and control NiTi stent implantation in bilateral CCAs and SCAs of beagles (indicated by red markings and arrows). Baseline DSA performed immediately after implantation confirmed successful deployment of both stent types at both sites, with optimal wall apposition and unobstructed blood flow. For the Mo stent, 3D-CTA at 90 days revealed structural distortion in the SCA but not in the CCA. By 180 days, DSA demonstrated progressive distortion in the CCA and re-coiling of the stent in the SCA. After 360 days, DSA showed further aggravation of the distortion in the CCA, while the previously re-coiled stent in the SCA also exhibited noticeable deformation. No significant thrombosis or in-stent stenosis was observed throughout the 360-day follow-up, and no apparent degradation of the Mo stent was detected. In contrast, the control NiTi stent group exhibited no signs of deformation or flow disturbance at any time point. Mo, molybdenum; NiTi, nickel–titanium; CCA, common carotid artery; SCA, subclavian artery; DSA, digital subtraction angiography; 3D-CTA, three-dimensional computed tomography angiography.

**Table 1.**
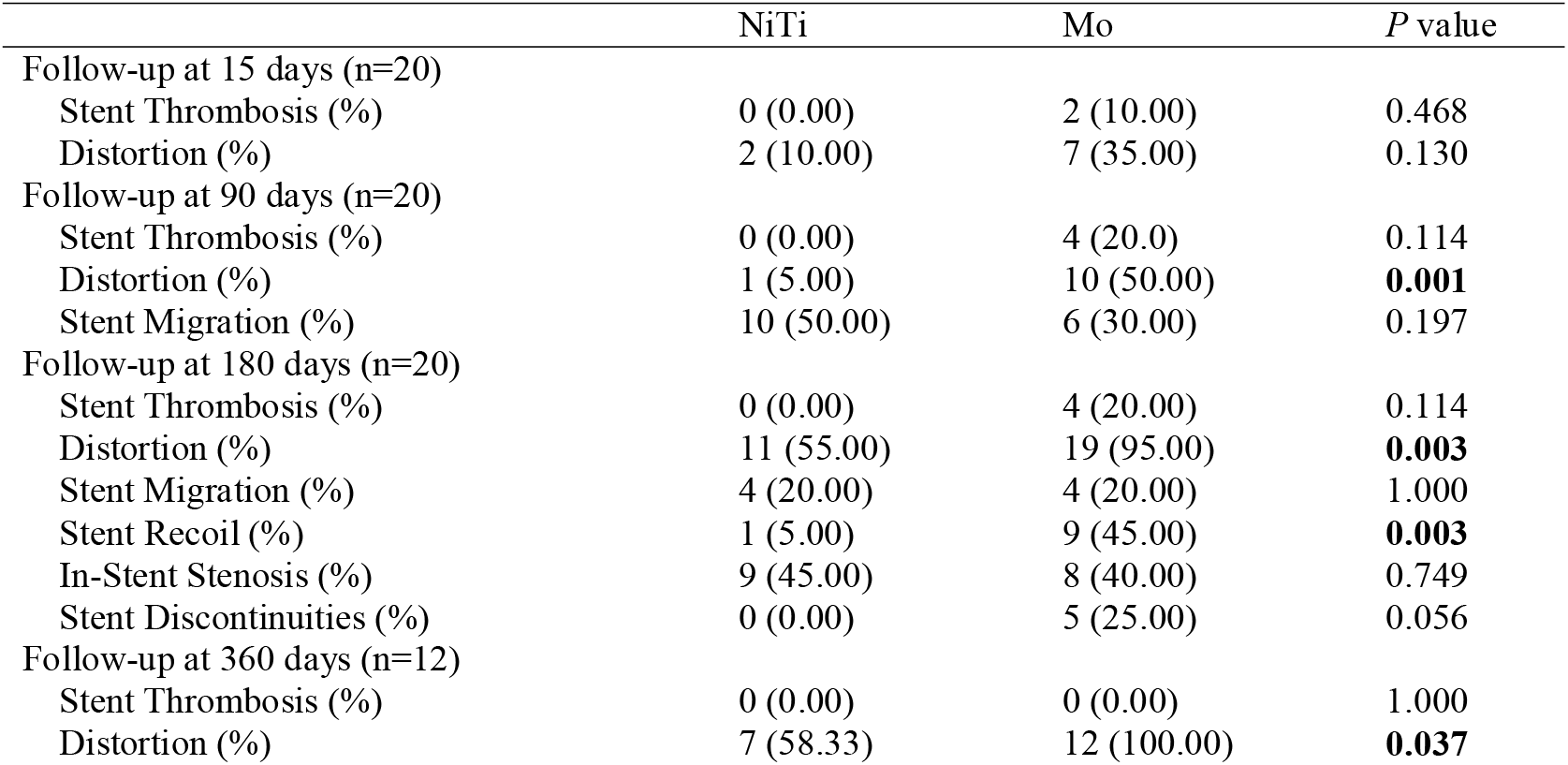
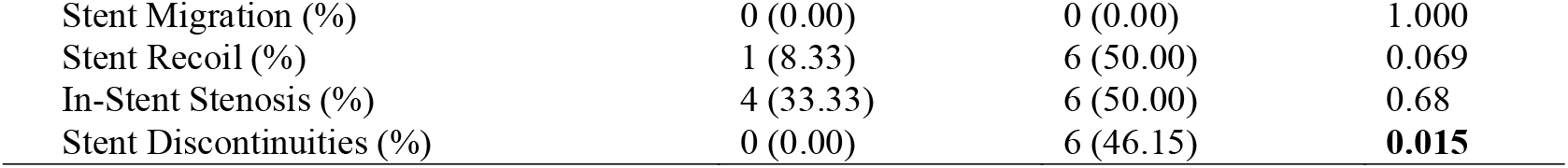
The structural changes and thrombosis events based on imaging results.

|  | NiTi | Mo | *P* value |
|---|---|---|---|
| Follow-up at 15 days (n=20) |  |  |  |
| Stent Thrombosis (%) | 0 (0.00) | 2 (10.00) | 0.468 |
| Distortion (%) | 2 (10.00) | 7 (35.00) | 0.130 |
| Follow-up at 90 days (n=20) |  |  |  |
| Stent Thrombosis (%) | 0 (0.00) | 4 (20.0) | 0.114 |
| Distortion (%) | 1 (5.00) | 10 (50.00) | **0.001** |
| Stent Migration (%) | 10 (50.00) | 6 (30.00) | 0.197 |
| Follow-up at 180 days (n=20) |  |  |  |
| Stent Thrombosis (%) | 0 (0.00) | 4 (20.00) | 0.114 |
| Distortion (%) | 11 (55.00) | 19 (95.00) | **0.003** |
| Stent Migration (%) | 4 (20.00) | 4 (20.00) | 1.000 |
| Stent Recoil (%) | 1 (5.00) | 9 (45.00) | **0.003** |
| In-Stent Stenosis (%) | 9 (45.00) | 8 (40.00) | 0.749 |
| Stent Discontinuities (%) | 0 (0.00) | 5 (25.00) | 0.056 |
| Follow-up at 360 days (n=12) |  |  |  |
| Stent Thrombosis (%) | 0 (0.00) | 0 (0.00) | 1.000 |
| Distortion (%) | 7 (58.33) | 12 (100.00) | **0.037** |
| Stent Migration (%) | 0 (0.00) | 0 (0.00) | 1.000 |
| Stent Recoil (%) | 1 (8.33) | 6 (50.00) | 0.069 |
| In-Stent Stenosis (%) | 4 (33.33) | 6 (50.00) | 0.68 |
| Stent Discontinuities (%) | 0 (0.00) | 6 (46.15) | **0.015** |

### 3.4 In vivo systemic toxicity and biocompatibility assessment

Histopathological analysis at 6 and 12 months post-implantation revealed no evidence of Mo accumulation or Mo-associated pathological lesions in the heart, liver, spleen, lungs, kidneys, or brain (Figure 3A, B). Immunohistochemistry and immunofluorescence analyses of the frontal, parietal, and occipital lobes showed no detectable abnormalities in neurons, astrocytes, microglia, or myelin structures (Figure 3B, S1). Systemic Mo was evaluated by measuring Mo concentrations in serum (sMo), urine (uMo), feces (fMo), along with serum uric acid (sUA), at baseline and multiple post-implantation time points (14, 30, 90, 180, and 360 days; Figure 3C). Serum Mo levels were significantly elevated relative to baseline at all post-implantation time points, with a gradual increase from 14 to 180 days and a more pronounced elevation at 360 days. In contrast, urinary Mo levels showed no significant difference from baseline at any time point. Although a transient rise was observed at 14 days, levels declined thereafter and remained stable through 90 to 180 days. A slight elevation in individual animals at 360 days did not result in a significant change in the group median. Fecal Mo levels remained stable throughout the study period, with no significant changes from baseline. To further assess systemic safety, hematological and biochemical parameters were monitored over 12 months (Figure 3D). The sUA levels were transiently elevated at 14 days but gradually declined afterwards. Hematological indices, including hemoglobin (HGB), white blood cells (WBC), and platelets (PLT), generally fluctuated within the normal reference ranges (indicated by the grey shaded areas), although transient elevations in WBC and PLT were observed at the early stage, consistent with an acute inflammatory response that resolved over time. Liver function markers—alanine aminotransferase (ALT), aspartate aminotransferase (AST), and alkaline phosphatase (ALP)—showed occasional statistically significant differences compared with baseline but remained largely within physiological limits. Similarly, renal function indicators, including creatinine (Cr) and blood urea nitrogen (BUN), showed no evidence of renal toxicity. Additional biochemical data are provided in Supplementary Figure S2.

**Figure 3.**
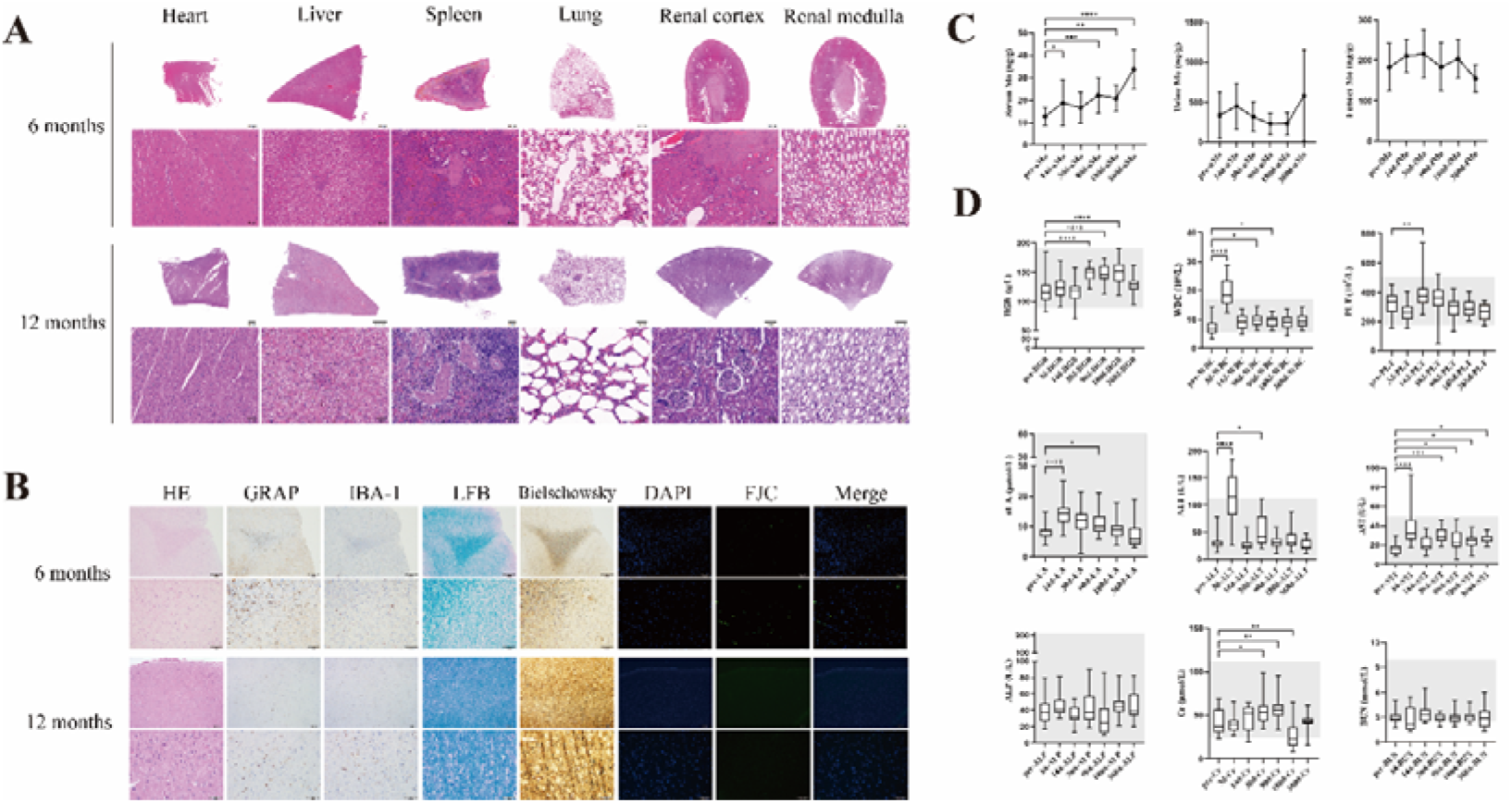
In vivo histologic staining analysis and laboratory test results. (A) H&E staining of heart, liver, spleen, lungs, renal cortex, and renal medulla at 6 and 12 months (0.8x and 20x magnification). (B) Immunohistochemistry and immunofluorescence staining of the parietal lobe at 6 months and 12 months (0.8x and 20x magnification). H&E staining revealed intact neuronal architecture with no signs of inflammation or necrosis. GRAP marked astrocyte activation; no significant hypertrophy or gliosis observed. IBA-1 indicated microglial activation status; minimal to no reactive microgliosis was detected. LFB demonstrated preserved myelin sheaths, with no evidence of demyelination. Bielschowsky staining visualized axonal integrity; axons appeared continuous and well-preserved without degeneration. DAPI: Nuclear counterstain showing normal nuclear morphology. FJC was used to detect degenerating neurons. Individual neurons with positive staining were observed. Based on H&E staining findings, these neurons exhibited normal structural features without degenerative changes. (C) Temporal profiles of molybdenum concentration in serum (sMo), urine (uMo), and feces (fMo), and (D) indicates hematological and biochemical parameters over a 12-month follow-up period, presented as box plots indicating the median, interquartile range, and minimum–maximum values. Normal reference ranges were indicated by the grey shaded areas. Statistical significance is denoted as \**p* < 0.05, ** *p* < 0.01, *** *p* < 0.001, \*\*\*\**p* < 0.0001 versus preoperative values; ns, not significant. Abbreviation: ALP, Alkaline Phosphatase; ALT, Alanine Aminotransferase; AST, Aspartate Aminotransferase; BUN, Blood Urea Nitrogen; Cr, Creatinine; DAPI, 4’,6-diamidino-2-phenylindole; FJC, Fluoro-Jade C; GRAP, Glial Fibrillary Acidic Protein; H&E, Hematoxylin-Eosin; HGB, Hemoglobin; IBA-1, Ionized Calcium-Binding Adapter Molecule 1; LFB, Luxol Fast Blue; PLT, Platelet count; UA, Uric Acid; WBC, White Blood Cell count.

### 3.5 In Vivo Mechanism and Degradation Behaviors Evaluation

H&E and Masson’s trichrome staining (Figure 4A, S3) at 6 and 12 months post-implantation demonstrated pronounced structural distortion of the Mo stents, accompanied by varying degrees of in-stent stenosis. Histological analysis revealed smooth muscle cell (SMC) proliferation and minor hemosiderin deposition within the vessel lumen. Furthermore, marked SMC and collagen fiber proliferation surrounding the Mo struts, along with scattered macrophage infiltration, were observed, indicating a localized inflammatory and remodeling response. In contrast, the NiTi control stents maintained their structural integrity without significant in-stent stenosis, showing only minimal macrophage presence and mild peristrut collagen deposition. Micro-CT analysis (Figure 4B) further characterized the temporal structural evolution of the Mo stent. At 6 months, the CCA stent largely preserved its tubular architecture, whereas the SCA stent exhibited early discontinuity and deformation. By 12 months, both stents showed pronounced structural distortion and segmental discontinuity, with evident recoiling, particularly in the SCA, suggesting accelerated degradation under higher mechanical loading conditions. Sites of discontinuity or severe deformation are indicated by red arrows.

**Figure 4.**
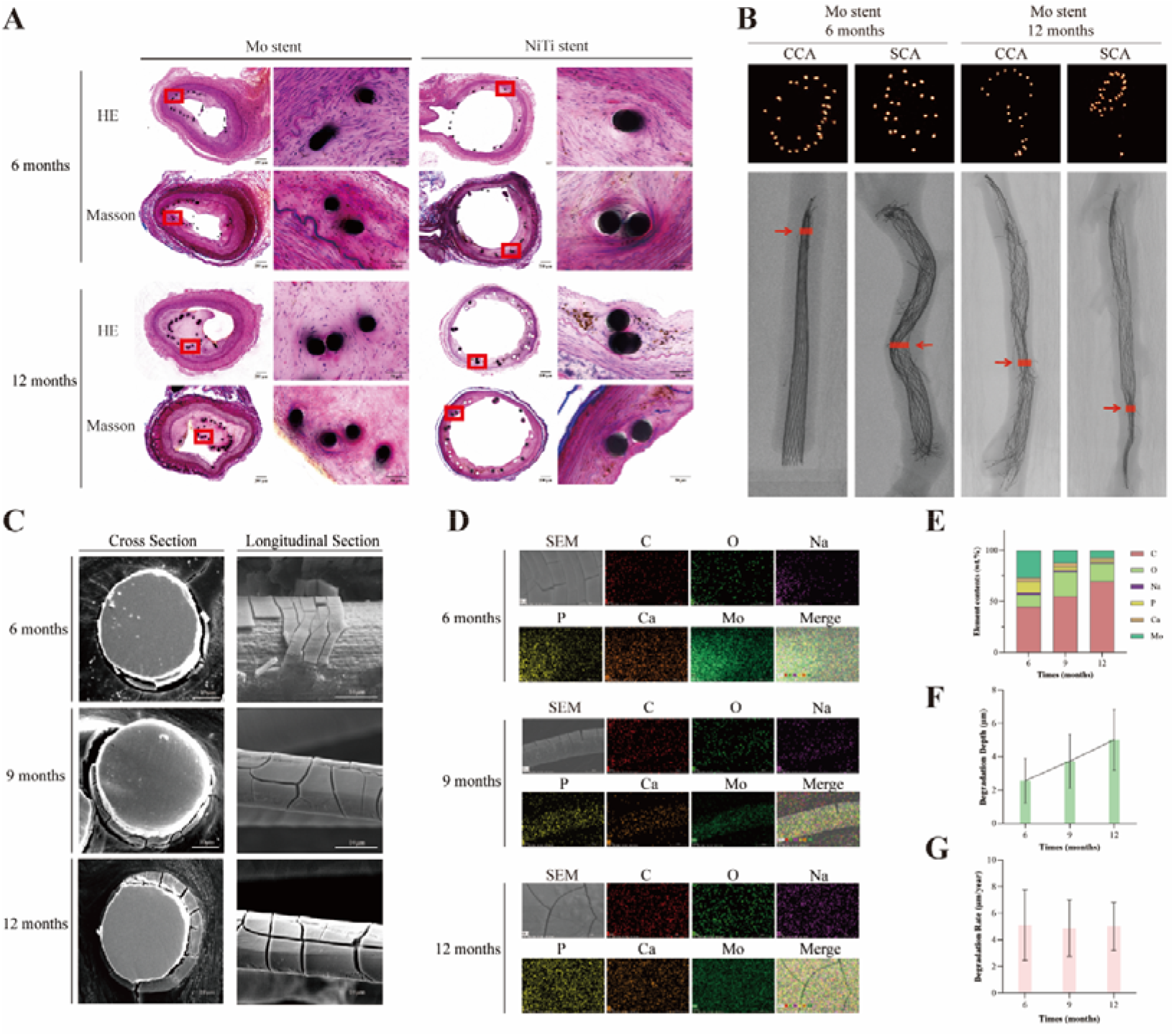
(A) The H&E and Masson’s trichrome staining of the Mo stents and NiTi control stents at 6 and 12 months post-implantation (4x and 40x magnification). (B) Micro-CT images of Mo stents implanted in the CCA and SCA at 6 and 12 months. The red arrows indicate the locations of cross-sectional images. (C) SEM image, EDS (D) elemental maps, and elemental composition (E) of the Mo struct at 6, 9, and 12 months. Degradation depth (F) and rate (G) of the Mo stent at 6, 9, and 12 months. Abbreviation: CCA, Common Carotid Artery; CT, Computed Tomography; EDS, Energy Dispersive Spectroscopy; H&E, Hematoxylin-Eosin; NiTi, Nickel-Titanium; SCA, Subclavian Artery; SEM, Scanning Electron Microscopy.

To elucidate the underlying degradation mechanisms, morphological and elemental analyses were conducted on retrieved stents at 6, 9, and 12 months. SEM images (Figure 4C) revealed a progressive reduction in strut thickness, consistent with relatively uniform corrosion. Longitudinal sections showed the presence of surface cracks, in agreement with in vitro immersion findings. However, the element analysis showed several differences. Elemental characterization by EDS (Figure 4D, E) identified, in addition to Mo, prominent signals of C, O, Na, P, and Ca. The enrichment of Ca and P suggested the formation of a calcium phosphate layer via biomineralization on the degrading surface. The surface layer is primarily composed of C and O. The elevated O content indicated an oxide layer, while the substantial C signal may reflect the presence of Mo carbonate species and/or adsorbed organic components from the physiological milieu. Collectively, these results indicate that in vivo degradation of Mo stents proceeds through oxidation followed by transformation into stable inorganic and organic-associated compounds.

Quantitative analysis of degradation kinetics (Figure 4F, G), based on cross-sectional measurements (Figure 4C), demonstrated a near-linear increase in degradation depth over time, corresponding to an estimated degradation rate of approximately 4.88–5.12 μm/year.

## 4. DISCUSSION

This study presents the first comprehensive one-year in vivo characterization of a braided pure Mo stent fabricated from 30.48 μm wire, validated in bilateral CCA and SCA implantation in Beagle dogs. The Mo stents exhibited favorable biocompatibility and a distinct degradation profile, with a slower in vivo degradation rate compared to in vitro conditions. Notably, time-dependent structural distortion was observed, particularly in the CCA and SCA, underscoring the critical interplay between material degradation and biomechanical loading in dynamic vascular environments.

The mechanical performance of the Mo stents revealed a distinct disparity between static in vitro testing and dynamic in vivo conditions. Although the stents exhibited superior radial strength (0.29 N) and a high breakdown potential (1840 mV) in vitro—suggesting robust corrosion resistance—they developed progressive structural distortion in vivo, beginning at 90 days. Notably, this deformation was more pronounced in the SCA than in the CCA, suggesting increased susceptibility to extrinsic mechanical loading. Both implantation sites more closely reflect clinically relevant vascular environments than the relatively stable abdominal aorta. We postulate that complex three-dimensional arterial deformations induced by neck-limb motion, particularly axial compression and bending, are the primary drivers of this phenomenon [26,27]. This aligns with findings by Xu et al., who demonstrated that dynamic vascular bending induces alternating stresses threefold higher than those from vascular pulsation[28]. Under such conditions, stress concentration at braided wire interconnections likely promotes localized mechanical failure, leading to structural discontinuity. This “extra-physiological” stress environment may accelerate fatigue crack initiation and propagation, ultimately resulting in recoiling and loss of structural integrity. Collectively, these findings highlight the necessity for next-generation bioabsorbable stents to incorporate enhanced fatigue resistance and mechanical adaptability to accommodate the complex loading conditions encountered in clinically relevant implantation sites.

The degradation behavior of the Mo stents further elucidates the difference between degradation and mechanically driven failure. At the time when structural deformation was first observed, the degree of cross-sectional loss of the struts remained insufficient to account for the loss of structural integrity based on material thinning alone. Accordingly, the observed failure is more consistent with the accumulation of mechanical fatigue under cyclic multiaxial loading, rather than premature chemically driven dissolution. This perspective is supported by previous studies. Bergström et al. indicated that under cyclic loading conditions, mechanical fatigue accumulation, rather than chemical dissolution, constitutes the dominant failure mechanism[29]. Similarly, Wildemann et al. demonstrated that fatigue accumulation under multiaxial loading is the primary determinant of progressive material degradation and eventual failure[30]. Together, these findings indicate a mechanistic decoupling between chemical degradation and structural deformation. This distinction is critical for the preclinical evaluation of biodegradable metallic stents and underscores the necessity of anatomically and biomechanically representative testing models that more accurately reproduce the complex loading conditions of target implantation sites.

Due to stent distortion and extensive tissue integration following long-term implantation, conventional degradation assessment via micro-CT or mass loss was unfeasible. Consequently, the degradation rate was evaluated by measuring the Mo wire diameter on unstained cross-sectional hard tissue sections using SEM. This method provided a direct assessment of structural reduction. Interestingly, the in vivo degradation rate was faster than that observed during the stable phase of in vitro degradation (approximately 4.88-5.12 μm/year vs. 3.75–3.98 μm/year). This discrepancy likely reflects the complex and reactive in vivo microenvironment. Upon implantation, the foreign body response activates macrophages and other inflammatory cells, which release protons and hydrolytic enzymes, thereby acidifying the peri-implant milieu and potentially accelerating Mo dissolution[31]. Concurrently, reactive oxygen species (e.g., H□O□, •OH) generated via enzymatic systems such as myeloperoxidase and NADPH oxidase can disrupt the passive Mo oxide layer and promote the formation of soluble high-valence Mo species[32]. In addition, continuous blood flow ensures sustained oxygen and electrolyte exchange, preventing ionic saturation and oxygen depletion that are often present in static in vitro systems. Although pulsatile hemodynamic forces and plasma protein interactions may further influence degradation, the stent struts became progressively endothelialized during implantation, which likely limited direct interaction with circulating proteins and reduced their contribution as dominant degradation drivers[33]. However, the observed in vivo degradation kinetics are compatible with the therapeutic timeline of intracranial aneurysms, most of which achieve complete occlusion within approximately 14 months [34]. In contrast, the current stents exhibit approximately 50% degradation at around 36 months. Future optimization should therefore aim to balance mechanical support during the critical healing phase with an appropriately accelerated and controllable degradation profile.

The EDS analysis of the degradation products revealed signals for Ca and P. This finding is likely analogous to the murine study by Sikora et al.[14], where compounds such as calcium molybdate, calcium phosphate, and calcium oxalate were identified on the degradation surface. These compounds likely contribute to the formation of a biomineralized surface layer on the Mo stent. In addition, the increased O and C signals observed in the EDS maps are consistent with the formation of Mo(III/IV) oxide species and the adsorption of plasma-derived organic components at the metal–tissue interface[21,22]. Collectively, these findings suggest that in vivo degradation of Mo proceeds through a coupled oxidation–biomineralization process rather than uniform electrochemical dissolution.

During endothelialization, Mo stents exhibited surface SMCs proliferation and collagen deposition on the luminal surface, accompanied by scattered macrophage infiltration. Notably, in-stent stenosis occurred predominantly at sites of structural distortion. Although the overall incidence of stent thrombosis and in-stent stenosis did not differ significantly between Mo and NiTi stents during follow-up, Mo stents demonstrated a higher trend toward these adverse events. These results differ from those reported in previous short-term in vivo studies[14,23,24]. However, similar phenomena have been observed in other biodegradable stents, such as poly-L-lactic acid stents. A network meta-analysis found that contemporary poly-L-lactic acid stents were associated with a significantly higher risk of stent thrombosis compared to drug-eluting stents [31]. Such adverse outcomes have been closely linked to stent deformation and subsequent endothelial dysfunction[35,36]. Specifically, early stent distortion may induce localized endothelial injury and trigger an acute inflammatory response, as evidenced by macrophage accumulation in histological analyses. This process, exacerbated by direct blood–strut interaction, may promote both thrombosis and neointimal hyperplasia leading to stenosis [37–40]. Although the hemocompatibility of Mo has been validated in previous studies, the elevated trend toward thrombosis and stenosis observed here is more likely attributable to mechanical deformation rather than intrinsic material incompatibility. Collectively, these results highlight the need for further optimization of Mo stent design, with particular emphasis on improving structural stability as well as systematically evaluating antithrombotic performance and resistance to in-stent stenosis under physiologically relevant mechanical conditions.

The 12-month systemic safety profile represents an important and reassuring outcome. The transient elevation of sMo and sUA observed within the first 14 days post-implantation is mechanistically plausible. In the early phase of stent implantation, the exposure of Mo wires to circulating blood likely facilitates the rapid release of degradation products into the systemic circulation. Mo serves as an essential cofactor for xanthine oxidoreductase (XOR), the enzyme responsible for the final steps of purine catabolism into sUA[21]. Accordingly, an acute increase in Mo ion availability may transiently enhance XOR activity, thereby increasing purine catabolism and sUA production. The subsequent normalization of sUA levels, coupled with the absence of a parallel increase in urinary Mo excretion, suggests that the released Mo ions are efficiently incorporated into physiological metabolic pathways and eliminated without systemic accumulation[25]. These findings highlight the importance of monitoring transient hyperuricemia in future translational studies, particularly in individuals with pre-existing hyperuricemic or gout-prone conditions. Histological examination of major organs (kidney, liver, spleen, heart, lungs, and brain) at 6 and 12 months revealed no Mo-associated pathological alterations, consistent with prior reports of Mo wire implantation in vivo[22,41]. Additionally, unlike the glomerular injury reported by Sikora-Jasinska et al. in mice[14], no such damage was found in the present beagle study. This discrepancy may reflect species-dependent differences in tolerance to Mo ion burden, with small animal models potentially exhibiting greater susceptibility than larger mammals. On this basis, Mo stents are unlikely to pose a significant renal toxicity risk in humans. Importantly, no evidence of neurotoxicity was detected using a multi-marker immunohistochemical assessment encompassing neuronal integrity, astrocytic and microglial activation, myelin structure, and axonal continuity. This is particularly relevant given the intended intracranial application of the device, and supports the favorable central nervous system biocompatibility of the Mo stent system.

Several limitations of the present study merit acknowledgment. First, the Beagle dog CCA/SCA model introduces mechanical demands that differ from those encountered in human intracranial arteries, particularly with respect to vessel diameter, wall compliance, and amplitude of non-pulsatile deformation. Second, the absence of an aneurysm model in the present study precludes direct evaluation of hemodynamic efficacy, including aneurysm occlusion rates, which represent the primary clinical endpoint of the intended application. Accordingly, the current findings are limited to device performance in a non-aneurysmal vascular environment. Third, while the 30.48 µm wire diameter outperforms most degradable materials and is comparable to clinically used braided stents, fatigue resistance remains a critical engineering constraint under long-term cyclic loading. Future studies should therefore focus on three main directions: (i) optimization of braiding parameters to achieve more uniform stress distribution within the stent architecture and mitigate localized fatigue; (ii) surface functionalization strategies, including pre-treatment via immersion in simulated body fluid, which has been demonstrated by Deng et al. to mitigate macrophage-induced inflammatory responses to Mo degradation products[42], as well as incorporation of drug-eluting coating to promote endothelialization and reduce thrombotic and restenotic risk during degradation[23], may accelerate luminal healing and further reduce thrombotic risk or in-stent stenosis during the degradation period; (iii) alloy optimization—such as Mo–Re systems—to enhance fatigue resistance while preserving biodegradability and biocompatibility[43].

## 5. CONCLUSION

This study demonstrates that braided Mo stents possess favorable mechanical properties and a degradation rate compatible with the therapeutic timeline of intracranial aneurysm treatment. In a one-year Beagle model, the stents exhibited good biocompatibility. However, structural failure associated with biomechanical fatigue, along with thrombotic risk and a trend toward in-stent stenosis, was observed. Collectively, Mo stents represent a promising class of biodegradable devices. These findings provide mechanistic and translational insights for next-generation design, highlighting the need to enhance fatigue resistance and optimize antithrombotic performance to ensure long-term structural integrity and clinical safety.

